# Mapping short reads, faithfully

**DOI:** 10.1101/2020.02.10.942599

**Authors:** Eduard Valera Zorita, Ruggero Cortini, Guillaume J. Filion

**Affiliations:** Center for Genomic Regulation (CRG), The Barcelona Institute of Science and Technology, Dr. Aiguader 88, Barcelona 08003, Spain; University Pompeu Fabra (UPF), Barcelona, Spain; Department of Biological Sciences, University of Toronto Scarborough, Toronto, ON, Canada

## Abstract

Mapping is the process of finding the original location of a DNA read in a reference sequence, typically a genome. Short read mappers are software tools used in most applications that involve high-throughput sequencing. As such, they must be continuously improved to keep up with increasing needs. Modern mappers rely on seeding heuristics, making them fast but inexact. For lack of a method to compute the reliability of their own output, mappers have so far used approximations of variable quality. Here we focus on faithfulness, the capacity to provide accurate mapping confidence, and we devise a strategy to map short reads faithfully. The key is to estimate the repetitiveness of the target reference, which is the dominant factor for the reliability of the mapping process. This approach highlights the existence of a class of reads that can be mapped with unprecedented confidence. We exploit this strategy in a prototype mapper that is competitive with state-of-the-art mappers BWA-MEM and Bowtie2, with the benefit of faithfulness. The software is open-source and available for download at https://github.com/gui11aume/mmp.

## 1 Introduction

High throughput DNA sequencing is now a well-established technology with countless applications in industry and medicine [1]. The Illumina short-read technology currently dominates the market of DNA sequencing over newer technologies that promise longer reads at the expense of higher error rates. Software and tools to process short-read sequencing data are therefore an important target for optimization [2].

In most analyses, short reads reads are mapped to a known reference, typically a genome, using a software tool known as a mapper. Mappers are complex algorithms that must solve approximate string-matching problems, where the reads and the reference do not match perfectly due to biological divergence or to measurement errors of the sequencing machinery. The traditional vision of mapping algorithms is to optimize speed and accuracy while maintaining a low memory footprint. From this perspective, a major breakthrough was the conception of the FM-index [3, 4], a data structure based on the Burrows-Wheeler transform [5] and the suffix array [6]. Modern short-read mappers such as BWA-MEM [7] and Bowtie2 [8] rely on specialized implementations of the FM-index for DNA sequences. More recently, new specialized algorithms such as RNA-seq mappers have further improved the speed and the accuracy for their specific applications [9].

However, speed, accuracy and memory usage are not the only metrics that matter for mapping algorithms. Equally important is an attribute called *faithfulness*.

Faithfulness comes into play for heuristic algorithms, *i.e.*, algorithms that are not guaranteed to return a correct result. In this context, a heuristic is *faithful* if it provides an accurate probability that the reported result is correct. Importantly, faithfulness is orthogonal to accuracy in the sense that a faithful algorithm is not necessarily accurate and *vice versa*.

The importance of faithfulness is recognized in the specifications of the SAM format [10], featuring a mandatory *mapping quality* field MAPQ, defined as “–10 ⋅ log_10_ *Pr*{mapping position is wrong}”. The concept was originally introduced in the MAQ mapper [11], but the proposed method cannot be carried to more modern mappers based on the FM-index.

More generally, there is no exact method to estimate these probabilities and most mappers rely on approximations that are valid only for some genomes and some sequencing technologies. The computations of mapping quality scores are usually undocumented, variable between mappers and versions, often spreading puzzlement and frustration in the community [12, 13]. At times, the standard is even disregarded as some mappers, like STAR [14], use the MAPQ field as a qualitative scale instead.

As a consequence, mapping quality is often neglected by users and developpers, even though it is critical in many applications. For instance, when calling *de novo* mutations, the standard procedure is to map sequencing reads to the genome of interest and identify mismatches as evidence for mutations. In this context, mapping errors cause artifacts that explain nonexistent mismatches, so the confidence in the final call is commensurate with the mapping quality. Inaccuracies of the mappers cause recurrent errors in cancer genomics [15, 16] where faithfulness would make a major difference.

Mapping quality and faithfulness are also important in multi-genome applications. For instance, when sequencing DNA from heterozyotes or chimeras, the confidence that a read is mapped to the correct location is in fact the confidence that the read is assigned to the correct genome. Applied to RNA-seq, for instance, mapping quality thus dictates the calls for mono-allelic expression. More generally, every analysis of this type depends on the confidence in the locations of the reads.

Improving the mapping process requires to understand why it is sometimes inaccurate in the first place. Mapping heuristics are usually based on a filtration strategy called *seeding*, whereby short exact matches between the read and the reference are used to extract a set of candidate locations. The downside of seeding is that errors in the read may cause the correct location to be filtered out, in turn the read to be mapped incorrectly. We recently developped a formal computational framework to compute the error rate of different seeding strategies used for mapping short reads [17].

Here we propose a method to estimate the unknown parameters of our previous model and we implement it in a mapper called MEM Mapper Prototype (MMP) that aims to be faithful. We use MMP to study the feasibility of this strategy and we highlight the additional benefits gained from faithfulness. In the process, we discover a class of reads that can be mapped with extremely high confidence and we show that faithfulness can be achieved without sacrificing speed or accuracy.

## 2 MATERIALS AND METHODS

### 2.1 Estimating the number of paralogs

The main source of uncertainty in the mapping is the existence of near-repetitive sequences, here referred to as paralogs. Paralogs are groups of sequences that share high levels of similarity, likely because they resulted from duplication or horizontal DNA transfer events. Paralogs are hard to map reliably because the read can be similar to some secondary candidates. At the same time, the exact probability that the read originated from one of the paralogs is difficult to quantify in practice. In this section we introduce a method to estimate the number of paralogs of a sequence. This estimate is crucial in the whole process of computing the mapping quality because the number of paralogs will be the main determinant of the quality.

We assume that the true location of the read is a sequence of the genome that has *N* ≥ 0 paralogs. For every nucleotide, the probability that a paralog differs from the target is a constant *µ* (this event is a mutation), and the probability that the read differs from the target is a constant *p* (this event is a sequencing error). We further assume that the genome has size *G* and that its composition is an equiprobable random mix of the four nucleotides.

For every nucleotide, the probability that the read differs from a given paralog is *λ* = (1 − *p*)*µ* + *p*(1 − *µ*/3). The probability that the paralog matches the last *L* nucleotides of the read is thus *a*_*L*_ = (1 − *λ*)^*L*^, and the probability that at least one of the the *N* paralogs matches them is 1 − (1 − *a*_*L*_)^*N*^. Finally, the probability that some paralog matches the last *L* nucleotides of the read, but none matches the last *L* + 1 is *ℓ*(*L|N*) = (1 − *a*_*L*+1_)^*N*^ − (1 − *a*_*L*_)^*N*^.

From the expression above, it is possible to estimate *N* by the maximum likelihood method. For this, we need to find the value of *N* that maximizes *ℓ*(*L|N*) for fixed *L*. This can be done analytically by solving *∂ℓ*/*∂N* = 0, yielding

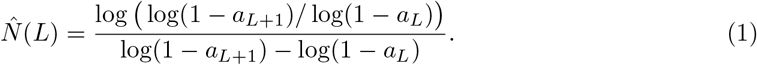

In practice, we can compute *L* with the backward search by extending the query until there are fewer than two hits in the genome (when there is only one hit, it is most likely the target). This is not exactly a maximum likelihood estimate because we should specifically account for the target as one of the hits. Also, *µ* is usually unknown, so we take a conservative approach and set *µ* to a value where the recall is lowest for reads in the range 50–150 bp (for MEM seeds we set *µ* = 0.06).

Finally, the formula is symmetric so we can obtain two estimates of *N*, one from each end of the read, and average them for more robustness. Note that expression (1) is a maximum likelihood estimate, but the final estimate of *N* is not, because of the approximations mentioned above.

### 2.2 Mapping quality

Based on the estimate of the number of paralogs, we define three different groups of reads. The mapping quality for each read group is computed differently because the dominant mapping uncertainty depends on the number of paralogs.

#### 2.2.1 Super quality

Super reads are defined as reads that satisfy the two following conditions: 20-mers extracted every 10 nucleotides all have a single hit in the genome (see the Results section and Figure 4), and the read is mapped without mismatch. Super reads are mapped to an incorrect location if:

1. the read contains *m* errors,
2. the target has exactly one paralog,
3. the paralog differs from the target exactly at the *m* locations of the errors,
4. the *m* incorrect nucleotides match the sequence of the paralog,
5. the *m* errors are in the odd-numbered 10-mers of the read.

The justification for these conditions are explained in more detail in Figure 5.

The last condition imposes *m* = 1 + [*L*/20] for a read of size *L*. The probability of condition 1 is denoted *α*. The probability of condition 2 is 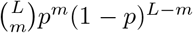, where *p* is the error rate of the sequencer. The probability of condition 3 is *µ*^*m*^(1 − *µ*)^*L−m*^, where *µ* is the probability that the paralog differs from the target. The probability of condition 4 is 1/3^*m*^ and the probability of condition 5 is 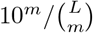

We need to multiply the probability of each condition together and divide them by the probability that a read is a *super read*, which is a fraction *β* of the probability that a read has no error. The mapping quality of super reads is finally computed as

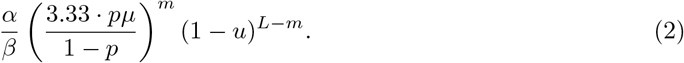

The parameter *µ* is set to *m/L* so as to maximize (2), and *α/β* was observed empirically to be close to 0.1.

#### 2.2.2 Normal quality

Reads with initial estimate *N* ≤ 20 are aligned to all the locations. The prior probability that the true location was not discovered at the seeding step can be calculated with `sesame` [17]. We can gain precision by computing the posterior probability given the number of mismatches (intuitively, a large number of mismatches is evidence that the read is mapped to an incorrect location). If the read is mapped to the correct location, the number of errors follows a binomial distribution where the parameter *p* is the error rate of the sequencer. If it is mapped to a paralog, the parameter is *λ* as defined above. A complication is that the distribution of errors is not binomial when *N >* 1 because the best hit is not chosen randomly among the *N* paralogs. We thus compute the posterior probablity using Bayes’ formula, but put an upper threshold equal to 1 on the log-ratio of the evidence. This means that many mismatches give strong support for an incorrect location, but few mismatches give only weak support for the correct location.

Assuming that the read is mapped with *x* mismatches, the probability that the location is incorrect is computed as

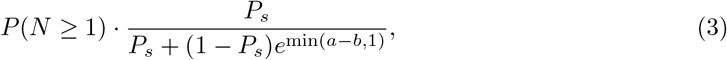

where *P*_*s*_ is the prior probability provided by `sesame`, where *a − b* = *x* ⋅ log(*p/λ*) + (*L − x*) ⋅ log((1 − *p*)/(1 − *λ*)), and where *P* (*N* ≥ 1) is a probability computed during the estimation of *N* (see below).

It was verified empirically that expression (3) is an overestimate when *x* = 0 (because here the upper threshold of 1 is too pessimistic). In this case, the estimate is replaced by a direct approach where we compute the probability that a paralog matches the read perfectly. More specifically, we compute the probability that the read has exactly one error and that a paralog matches the incorrect sequence. In this case, the probability that the location is incorrect is computed as

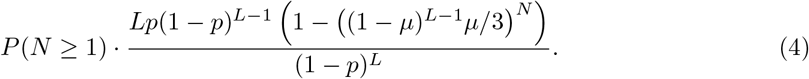

In both (3) and (4), *µ* is set to 0.06 as this gives a high proability that MEM seeding fails for reads of size 50–150.

#### 2.2.3 Low quality

Reads with initial estimate *N >* 20 are aligned only to the longest MEM seed. A seed of size *S* exists if there are *S* nucleotides without error or if there are *S* nucleotides with possibly many errors matching at least one of the *N* paralogs. In the first case, the target is discovered, in the second it is not. We also have to know whether the seed is flanked by errors or by the end of the read. The probability of the first case is

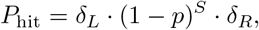

where *δ*_*L/R*_ = 1 if the seed is flanked by the end of the read and *δ*_*L/R*_ = *p* otherwise.

Considering only the most frequent case with a single error, the probability of the second case is

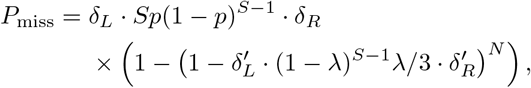

where *λ* is defined as above, and where 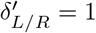 if the seed is flanked by the end of the read and 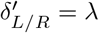 otherwise. Applying Bayes’ formula, the mapping quality is finally computed as

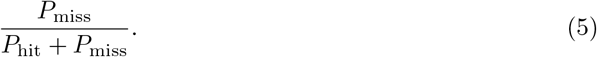

#### 2.2.4 Estimating *P* (*N ≥* 1)

This quantity appears in expressions (3) and (4). The estimate is based on the method shown in Figure 3, where we use the end of the read with the shortest query. We consider three cases: *i.* the target has a paralog, *ii.* only one end of the target has a paralog, and *iii.* the target is unique. Case *ii.* can occur when the read straddles the end of a transposon or other repeated sequence. We consider that *N* ≥ 1 only in case *i.* where the sequence has a full-length paralog.

We denote *m* the longest query size with two or more hits in the genome (see Figure 3) and we work the probability *P* (*m*) in the three cases above. The genome has size *G* and every nucleotide matches the query with probability 1*/*4 if it is unrelated to the target, and with probability *λ* if it is a paralog. For convenience, we also denote the probability of a random hit of size less than *m* as *ξ*(*m*) = 1 − 1/4^*m*^.

In case *i*., the observed value is *m* if there is a match of size *m* with the paralog at one end of the read, and a match of size ≥ *m* at the other end, and a match of size *< m* with the rest of the genome; or if the match with the paralog has length *< m* at both ends of the read, and there is a match of size *m* with the rest of the genome; or if there is a match of size *m* with the paralog at one end of the read, and a match of size ≥ *m* at the other end, and a match of size *m* with the rest of the genome. In case *i*., *P* (*m*) is thus

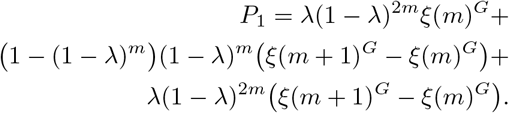

In case *ii.*, the observed value is *m* if there is a match of size ≥ *m* with the paralog at the duplicated end, and a match of size *m* with the rest of the genome at the other end; or if there is a match of size *< m* with the paralog at the duplicated end, and a match of size *m* with the rest of the genome at any end, and a match of size *≥ m* at the other end. In case *ii.*, *P* (*m*) is thus

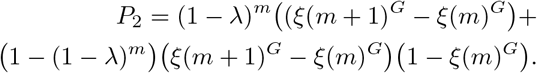

In case *iii.*, the observed value is *m* if there is a random match of size *m* with the genome at one end of the read, and a random match of size *≥ m* at the other end. In case *iii.*, *P* (*m*) is thus

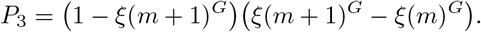

Finally, to compute the probability of case *i.* given the observed value of *m*, we use Bayes’ formula with prior odds in proportion 1: 1: 8 because we estimate that the probability that a sequence has exactly one paralog is approximately 1*/*10 (remember that those calculations are performed when there is evidence that *N ≤* 20).

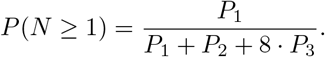

## 3 RESULTS

### 3.1 General design principle of a faithful mapper

Implementing a faithful mapper requires a theory to estimate the probability that the location of a read is wrong. We recently showed that most of the errors can be attributed to the seeding step and we developed a computational framework to estimate the error rate of different seeding schemes [17]. If we want those probabilities to correspond to the overall error rate of the mapper, it is important to verify all the candidate locations after the seeding step, otherwise neglecting some candidates would introduce some further mapping errors with unknown probability.

We thus opted to implement a mapper based on MEM seeds (where MEM stands for Maximal Exact Match) because this strategy produces a smaller candidate set than the alternatives. In order to optimize the design of the filtration step, we used the `sesame` library [17] to sketch the projected performance of MEM seeds (Figure 1).

**Figure 1:**
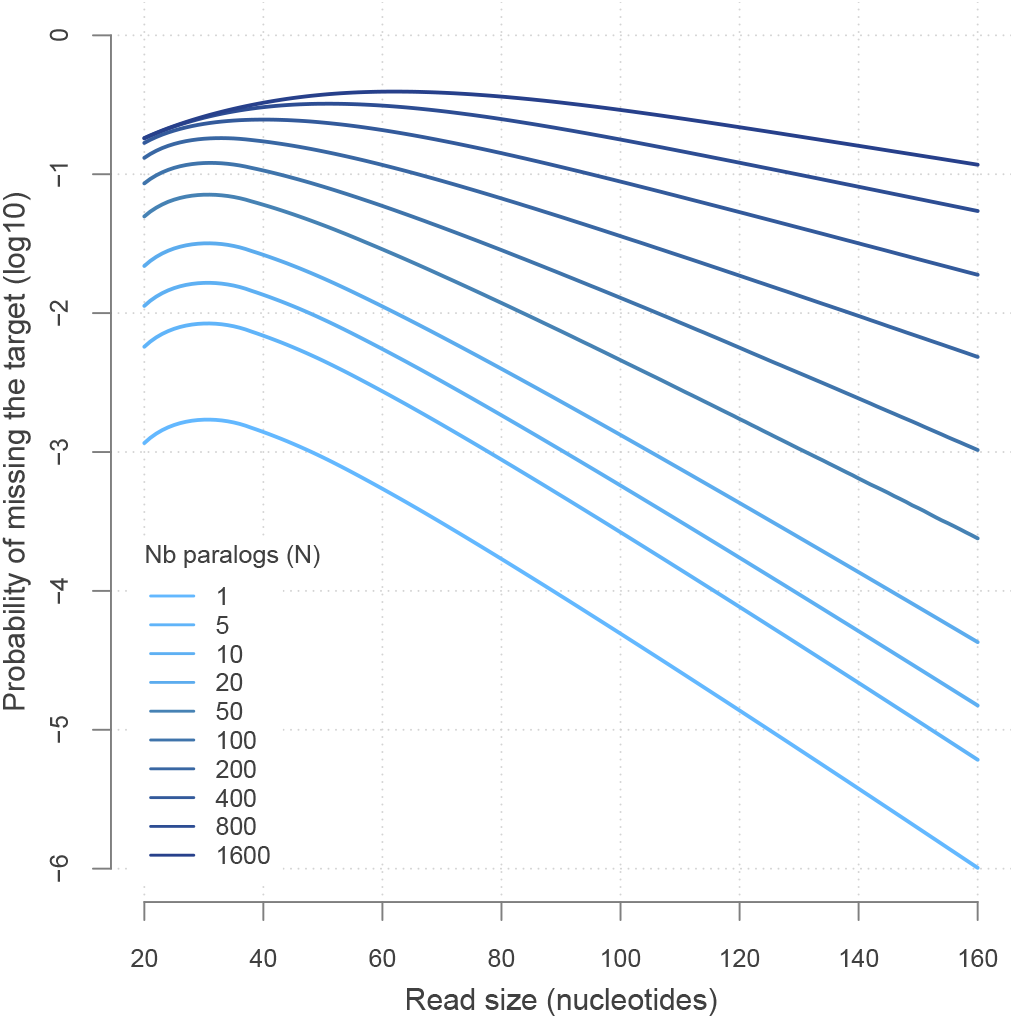
Projected performance of MEM seeds. The probability of filtering out the target is plotted against the size of the read (notice the log scale on the *y* axis). Computations are carried out with `sesame` for MEM seeds of minimum size 19, assuming an error rate of 1% and a divergence rate of 6% between paralogs (*i.e.* the probability that a given nucleotide of the target is different in a given paralog is 0.06). Each line represents the probabilities for a different number *N* of paralogs. Note that the asymptotic decay is approximately constant up to *N* = 20 and slows down for larger *N*.

The results reveal two essential insights: The first is that the asymptotic decay of the failure rate is approximately constant when the read maps to a location with *N* ≤ 20 paralogs. Beyond this, the asymptotic decay slows down and the recall decreases markedly. The second insight is that up to 100 nucleotides, the failure rate is always above 1 mapping error per 100,000 reads. In other words, there is no hope to map a read with very high confidence using MEM seeds if the target has a paralog.

These insights suggest that an efficient strategy is to use a fast mapping mode when the target has many paralogs because the read cannot be mapped with high confidence anyway. In contrast, when the number of paralogs is low, it is important to know whether the target is a unique sequence because this is the only way to map a read with high confidence. So after MEM seeding, our strategy is to quickly estimate the number of paralogs, find a decent candidate location as fast as possible and finally estimate how reliable this location is. In essence, we search the reads that can be mapped with high confidence instead of searching a high-confidence location for every read. A high-level summary of this strategy is shown in Figure 2.

**Figure 2:**
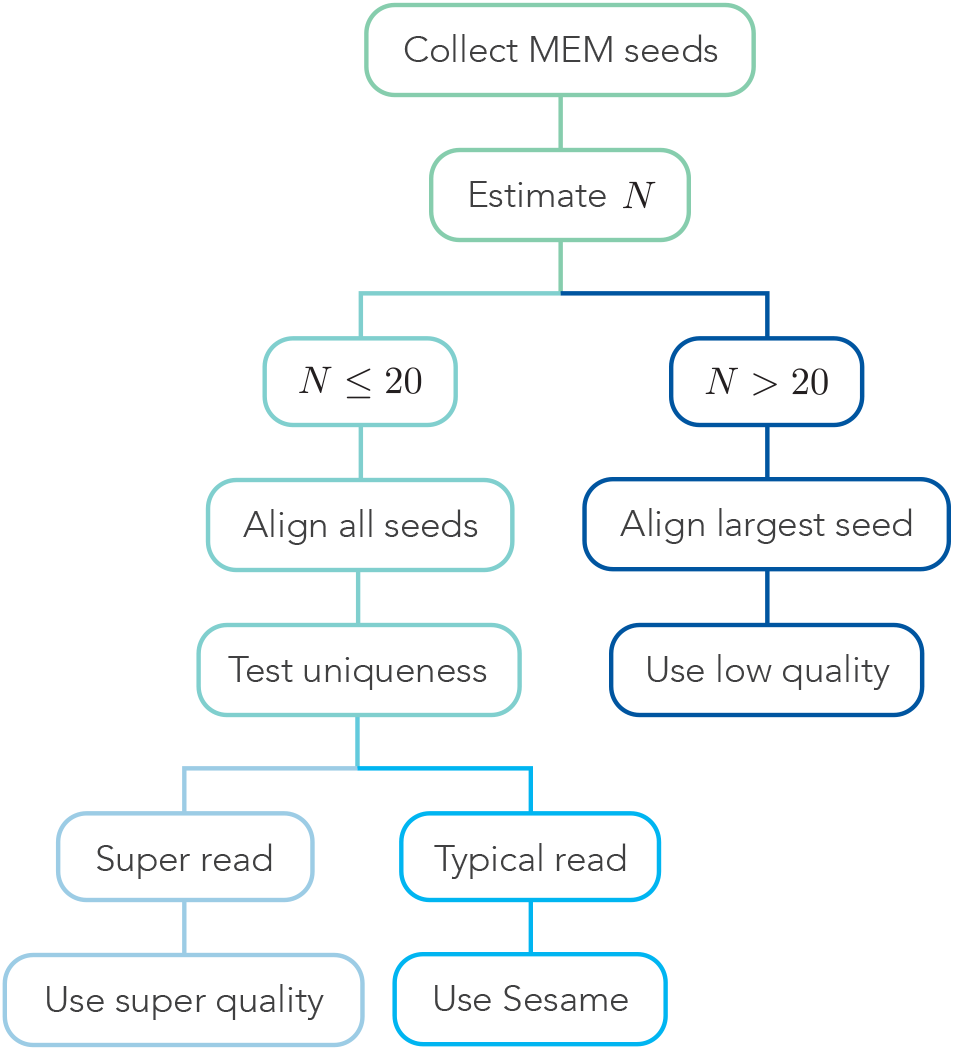
Proposed strategy for a faithful mapper. Each read is routed through one of three mapping pathways. First, all the MEM seeds are extracted (top). Then the number *N* of paralogs of the target is estimated. If *N* is 20 or less (left branch), all the candidates are tested with the Needleman-Wunsch algorithm. Once the best candidate is found, the probability that it is unique is estimated. If it is high and the alignment has no mismatch, mapping quality is computed using expression (2). If the best candidate is not unique or if the alignment has a mismatch, mapping quality is computed using expression (3) or (4). If *N* is higher than 20 (right branch), only the largest MEM seed is used and the mapping quality is computed using expression (5).

### 3.2 Estimating *N*

The key in the flow chart of Figure 2 is to dispatch the reads based on the number *N* of paralogs of the target, which is an indirect measure of the mappability of the read [18]. Estimating *N* must be fast so that this step does not become a bottleneck. To achieve this, we propose to use the seeding process itself. MEM seeds are best computed using the backward search [3], an algorithm that returns the number of hits in the genome as the query is extended backward. If the target has no or few paralogs (Figure 3a), the number of hits will quickly go below 2 (when there is only one hit, it is most likely the target itself). In contrast, if the target has many paralogs (Figure 3b), the number of hits will remain at 2 or above for many additional iterations.

**Figure 3:**
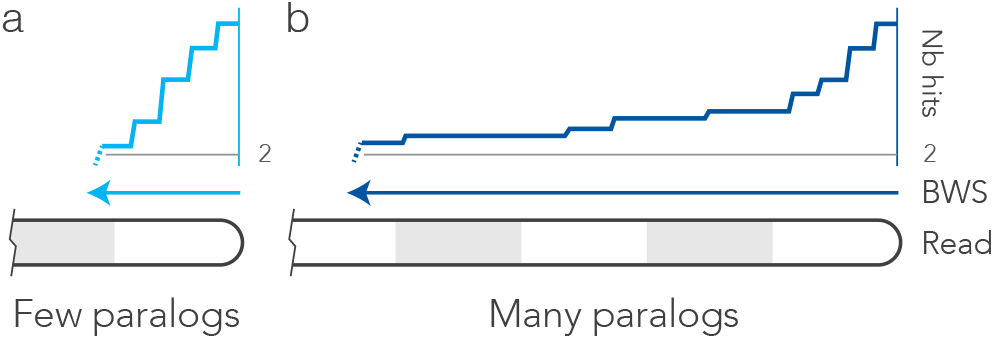
Estimating *N* with the backward search. **(a)** The target has no or few paralogs. It takes only a few iterations until the backward search (BWS) finds fewer than 2 hits in the genome. **(b)** The target has many paralogs. In this case it takes more iterations before the query has fewer than 2 hits. The number of iterations before hitting the threshold is used as a statistic to estimate *N*.

We can thus use the length of the largest query with 2 or more hits in the genome as a statistic to estimate *N* (see Materials and Methods).

### 3.3 Super reads

As mentioned above, our strategy to map reads with high confidence is to test whether the target is unique. Intuitively, a read that maps to a sequence without paralog in the genome is very likely to be mapped to the correct location.

We designed another estimation procedure based on the backward search, where this time we use the best candidate location as a query instead of the read. The principle, sketched in Figure 4a, is to perform the backward search on 20-mers and to test whether the genome contains a single hit. The rationale is that the total number of 20-mers is approximately 10^12^, so the chances of spurious random hits are very low, even in the largest known genomes.

**Figure 4:**
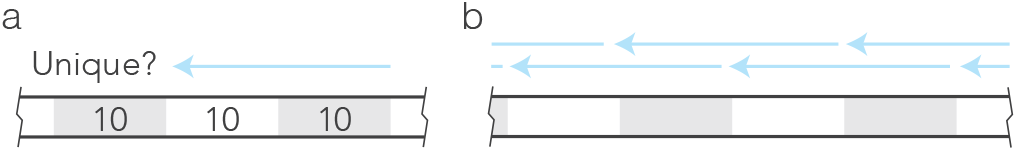
Test for uniqueness. **(a)** Basic principle. After the alignment stage, the sequence of the best hit is segmented in 20-mers. The uniqueness of a given 20-mer is tested with the backward search. **(b)** Complete test. The genomic location returned by the search process is considered to be unique if all the 20-mers overlapping by 10 nucleotides are unique.

For a complete test, the candidate location of the read is segmented in 20-mers overlapping by 10 nucleotides, as shown in Figure 4b. The sequence is considered unique if all the extracted 20-mers have a single hit in the genome.

While testing this procedure, we noticed that a class of reads were mapped with higher confidence than ever reported before. The *super reads*, as we called them, were those aligning without mismatch to a unique location.

The extremely high mapping quality of super reads comes from an unexpected property of the test depicted in Figure 4. To see why they can be mapped with such confidence, consider a read aligning without mismatch to an incorrect sequence of the genome. Observe that for this to happen, the read must contain one or more errors that are perfectly compensated in some paralog of the target. This can only happen if *i.* the target has a paralog, *ii.* the read contains an error, *iii.* this error matches the paralog and *iv.* the target and the paralog are otherwise identical.

But those conditions are not sufficient to make a read *super*. As shown in Figure 5a, the mapping location would not be considered unique because there would exist at least one 20-mer where the target and its dupliate are identical, yielding two hits. For a read of size 50, the paralog can be overlooked only if there are at least two errors in the read that match the paralog.

**Figure 5:**
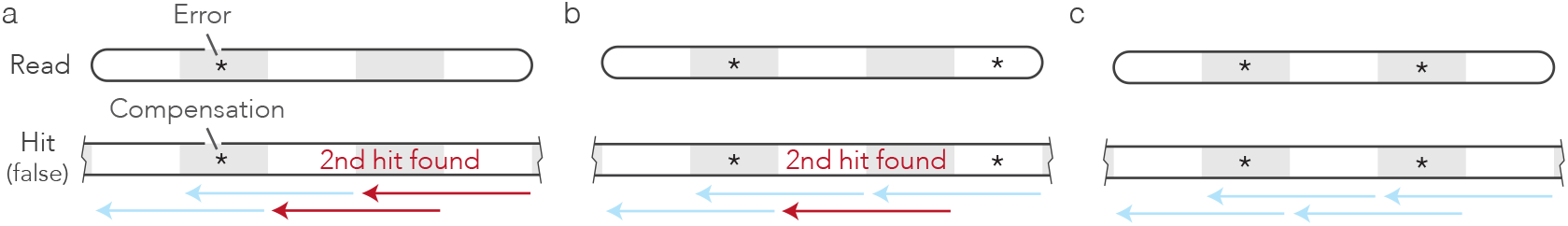
Properties of super reads. **(a)** Incorrect alignment with no mismatch. For a read to align perfectly to an incorrect sequence, it must contain an error (star in the read) and the incorrect nucleotide must match a paralog (star in the hit). This is not a super read because some extracted 20-mers are not unique (red arrows). **(b)** Incorrect alignment with several errors and no mismatch. Adding compensated errors is not always sufficient to make 20-mers unique: too much space between errors can leave 20-mers that are not unique, in which case the read is not super. **(c)** Incorrectly mapped super reads. If the compensated errors are in odd-numbered 10-mers (grey boxes), then all the extracted 20-mers are unique and the read is super. Those conditions are exceedingly rare, so super reads are mapped with high confidence.

But as shown in Figure 5b, this is still not sufficient for a wrongly mapped read to qualify as *super*. For this, the errors must be in every second 10-mer, as indicated in Figure 5c. In this configuration, the errors mask the presence of another hit on all the tested 20-mers and the incorrect target location is considered unique. There are other scenarios where a super read is mapped to an incorrect location, but they involve even more compensated errors so they can be neglected.

### 3.4 MEM Mapper Prototype

We implemented the mapping strategy presented in Figure 2 in a prototype mapper called MEM Mapper Prototype (MMP). MMP is implemented as a stand-alone C program including the code of `sesame` [17] and of `divsufsort` [19]. We used plain mapping algorithms to better highlight the benefits of building a mapper for faithfulness. The index consists of a standard FM-index of the genome and its reverse complement [the implementation is detailed in ref. 20]. In line with the GEM mapper [21], we also added an auxiliary lookup table storing the states of the bacwkard search for all possible 12-mers, allowing us to skip the first 12 iterations for every query. This lookup table has a fixed size of 256 MB. Sequence alignment is carried out using a version of the Needleman-Wunsch algorithm [22] with an option to abort if the score of the current best hit is exceeded (continuing the algorithm is useless to find the optimal location of the read). Mapping quality is computed as detailed in the Materials and Methods section.

The implementation otherwise follows the flow chart of Figure 2, with two differences. The first is that super reads can have 0, 1 or two mismatches, as long as all the extracted 20-mers have a single hit in the genome (the mapping quality is modified accordingly). The second difference is that once the best hit is found, *N* is estimated again using the genomic sequence instead of the read, and the higher of the two estimates is kept.

### 3.5 Benchmark

We set up a test bed using the genomes of *Drosophila melanogaster* (the fruit fly), *homo sapiens* (modern humans) and *Pinus taeda* (a north American species of pine). The high-level features of the genomes are summarized in Table 1. We included *Drosophila* and human as model organisms with high-quality genome assemblies, and pine as a non-model organism with a very complex genome and a draft assembly.

**Table 1:**
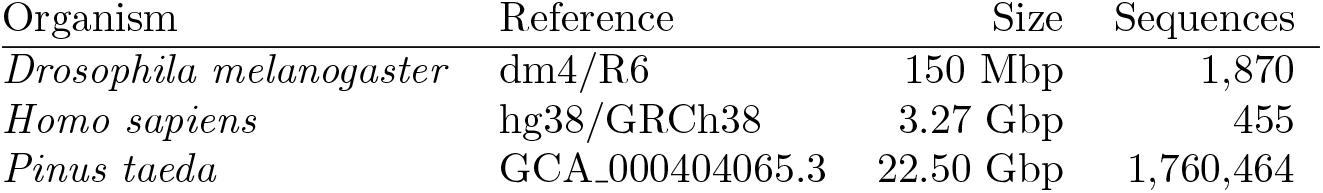
Features of the genomes used for benchmark.

The data sets used for the benchmark consist of 50 million sequences drawn uniformly at random from each genome. To simulate sequencing errors, nucleotides were randomly substituted with a fixed probability; there were no insertions or deletions. For each data set, the sequences have the same length (50, 75 or 100 nucleotides) and the same error rate (1%, 2%, 5% or 10%).

To set a baseline for comparison we used BWA-MEM and Bowtie2. The purpose of the benchmark is to test faithfulness as a viable mapping strategy, not to run a comprehensive survey of each mapper on the data set. Each mapper was thus used with its default parameters (but we passed the error rate to MMP, *e.g.*, `-e .02` for a 2% error rate). We used BWA-MEM version 0.7.9a-r786 [7], Bowtie2 version 2.3.5.1 [8], and MMP version 1.0. All the mappers were run in single thread.

The mapping experiments were run on a Hewlett Packard Z800 workstation with 64 bit Intel Xeon CPU X5675 at 3.07 GHz, with 64 GB of DDR3 RAM, 12 MB of cache, and running Linux Ubuntu 18.04.3. Indexing the pine genome with MMP had to be done on a machine with 512 GB of RAM because of the high memory footprint used by the library `divsufsort`.

We measured faithfulness by comparing observed versus claimed MAPQ score on simulated data. Figure 6 shows a series of scatter plots at 1% error rate on simulated reads. The area of the circles is proportional to the number of reads in the given category, and pink circles indicate that the observed mapping quality had to be imputed because all the reads were mapped correctly.

**Figure 6:**
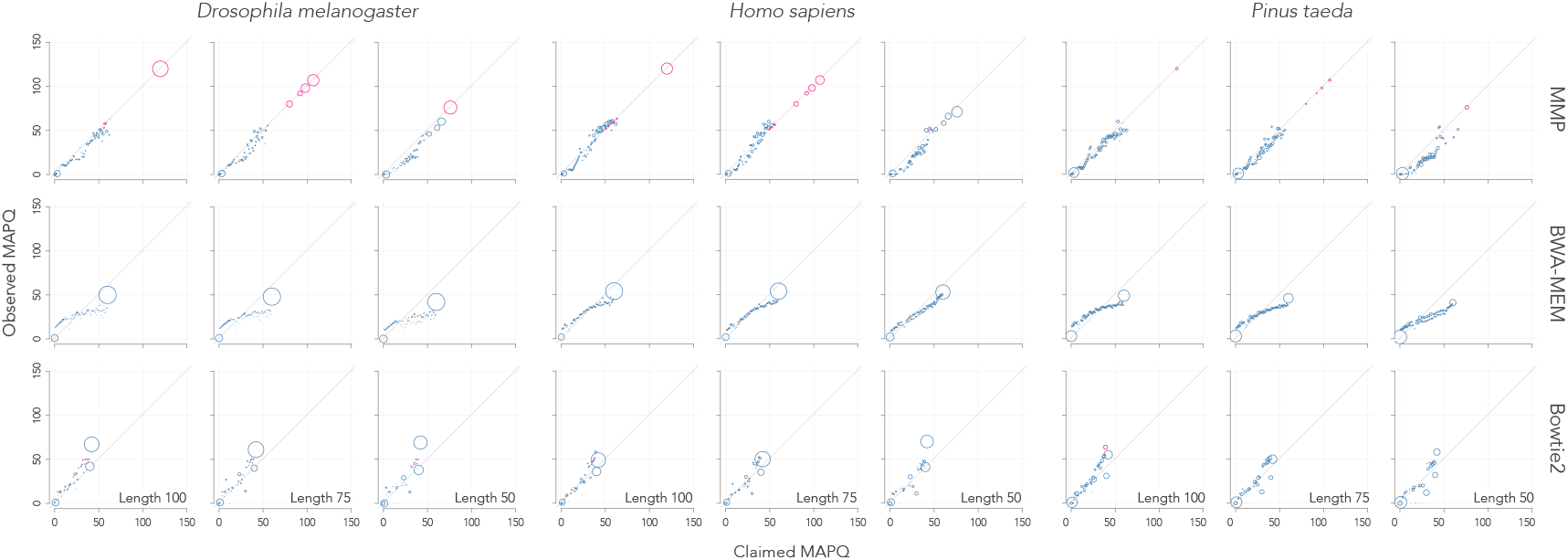
Faithfulness with 1% error rate. The observed mapping quality (MAPQ) is plotted against the value computed by MMP, BWA-MEM or Bowtie2. The data is showed for reads of size 100, 75 or 50, sampled from the genome of the fruit fly (*D. melanogaster*), of modern humans (*H. sapiens*) or of the pine (*P. taeda*). The size of the circle is proportional to the number of reads in the given category. Pink circles indicate that no mapping error was observed, so that the observed mapping quality was undefined. In this case one pseudo error was added if the expected number of errors was higher than 1 (we should observe errors but we did not), otherwise the observed mapping quality was set to the claimed mapping quality (we should not observe errors and we did not).

The faithfulness of MMP compares favorably to that of BWA-MEM and Bowtie2 because on average, the points lie closer to the diagonal. BWA-MEM tends to overestimate mapping quality in the high range (except for the human genome) whereas Bowtie2 tends to understimate it. Importantly, MMP explores a wider range of mapping qualities, allowing it to map reads with extremely high confidence. Those high-confidence reads with MAPQ above 60 are all super reads, and Figure 6 shows that in typical mapping conditions, their number can be very high (up to two thirds when mapping 100-mers in the *Drosophila* genome). Intuitively, the number of super reads goes down as the complexity of the genome increases.

Similar analyses for reads with higher error rates are shown in Supplementary Figures 5, 6 and 7. The faithfulness of MMP is equally good at 2% error rate, but it decreases substantially at 5% and 10% error rates, showing that our strategy is valid only within the scope of current short-read sequencing technologies.

**Figure 7:**
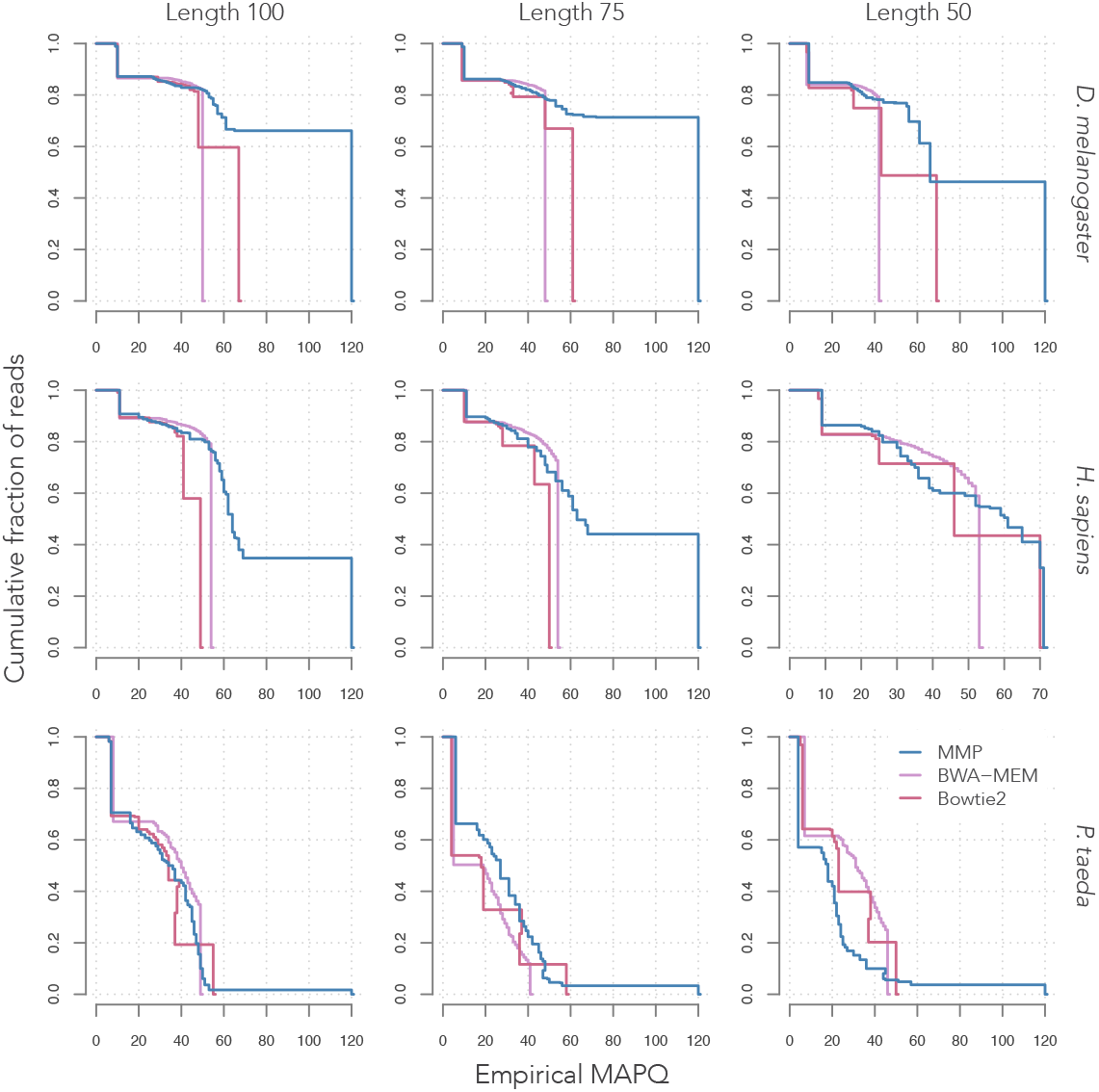
Mapping accuracy with 1% error rate. Plotted in each panel is the cumulative fraction of the reads above a certain mapping quality for MMP, BWA-MEM or Bowtie2. The score is computed empirically by sorting the reads on their MAPQ score and computing the average mapping quality of the reads in the top *x*% for all values of *x*. The data is showed for reads of size 100, 75 or 50, sampled from the genome of the fruit fly (*D. melanogaster*), of modern humans (*H. sapiens*) or of the pine (*P. taeda*).

It is important to evaluate the impact of faithfulness in terms of accuracy, speed and memory footprint. Figure 7 shows that the accuracy of MMP is competitive with that of BWA-MEM and Bowtie2 (recall that the mappers were used with default parameters). On the *Drosophila* genome, MMP is more accurate at most confidence levels, whereas there are more variations on the human and the pine genomes. The worst case for MMP is that of 50-mers mapped in the pine genome, perhaps because a MEM-only seeding strategy is not competitive in such cases. At 2% error rate, MMP shows a neat advantage in accuracy in most conditions (Supplementary Figure 2) but at 5% and 10% error rates, *i.e.* outside the realm of current short read technologies, the performance of MMP is more variable (Supplementary Figures 3 and 4). Those results show that optimizing for faithfulness can resut in good performance for accuracy.

In terms of speed, MMP is 2–4 faster than BWA-MEM and Bowtie2 on the present data set (Figure 8a), with a memory footprint that is approximately 2–3 times higher (Figure 8b). Part of the speed-up is due to the implementation and the larger stress on memory, but the strategy highlighted in Figure 2 aims to reduce the number of alignments when reads cannot be mapped accurately. For instance, MMP spends 8.3% of the time on incorrectly mapped reads, versus 29.2% and 12.5% for BWA-MEM and Bowtie2, respectively (100 nucleotide reads in the human genome, data not shown). At 2% error rate, MMP remains the faster mapper, but at 5% and 10% error rates, Bowtie2 can be somewheat faster (Supplementary Figure 1). However, at such high error rates, the mappers have poor accuracy and the running time is rather a reflection of how fast they “give up” on mapping the reads.

**Figure 8:**
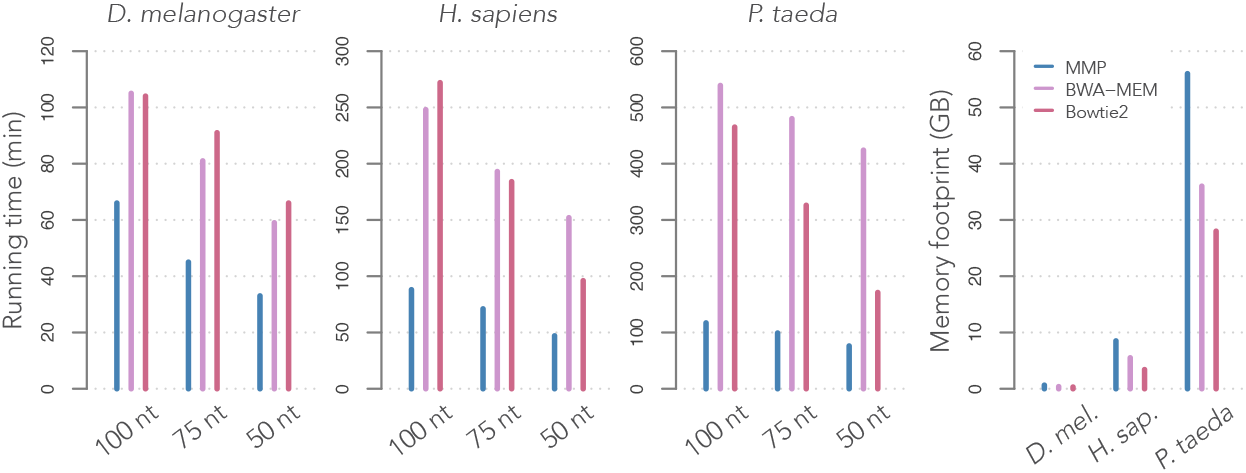
Resource usage at 1% error rate. **(a)** Speed. The running time is plotted for MMP, BWA-MEM or Bowtie2. The data is showed for reads of size 100, 75 or 50, sampled from the genome of the fruit fly (*D. melanogaster*), of modern humans (*H. sapiens*) or of the pine (*P. taeda*). **(b)** Memory usage. The peak memory footprint is plotted for the different mappers with reads from different genomes (the memory footprint is the same for all read sizes).

Overall, Figures 7 and 8 show that the investment in faithfulness is paid back in several ways: First, it gives access to very high confidence levels. Second, it auto-tunes accuracy by better separating low from high-confidence reads. Third, it allows the mapper to save time on low-confidence reads. Taken together, our results thus show that faithful mapping is not only a viable strategy: it also opens opportunities for further optimizations that can make the mapping process more efficient.

## 4 DISCUSSION

Here we designed and implemented a strategy to map short reads faithfully. The principles are based on key insight gained from models to compute seeding probabilities [17], showing that MEM seeds are efficient only when the target has few paralogs (Figure 1). This motivates the design of a strategy where we first estimate the number of paralogs of the target (Figure 3) to dispatch reads to different mapping subroutines (Figure 2). This amounts to allocating more resources to the most mappable reads [18]. This process revealed the existence of super reads, *i.e.*, reads that can be mapped with extremely high confidence (Figures 4 and 5), together setting the basis of an efficient mapping strategy. We wrote a mapper based on this strategy and we showed that it is competitive with BWA-MEM and Bowtie2 (Figures 6, 7 and 8). Importantly, the mapping process itself relies only on standard algorithm, so all the benefits of the mapper come from faithfulness. Overall, this demonstrates that improving faithfulness is a fruitful strategy to improve short-read mappers.

### 4.1 Estimating MAPQ

Our previous work indicated that seeding is the critical step to estimating mapping quality [17]. Computing seeding probabilities requires to know the error rate of the sequencer, the number *N* of paralogs of the target and their sequence divergence. The error rate may be known with reasonable accuracy, but *N* is typically less reliable, which is one of the major difficulties with the strategy developed here, and also the main reason why the mapping quality of MMP is sometimes inaccurate (Figure 6).

`sesame` computes probabilities with high precision [17], but several practical considerations obstruct the estimation of *N*. The first is that estimating *N* must be fast. The method depicted in Figure 3 only requires a backward search at each end of the read, but it provides only two measurements and is thus noisy. The second practical consideration is that the left and right halves of the read may have different values of *N*. This is the case, for instance, when the read straddles a transposon, where the end in the transposon may have a high copy number and the other end may be unique. The final consideration is that the evolutionary model assumes that the paralogs drift away from each other at the same speed, which ignores their genealogy. The estimation thus provides a *proxy* estimate of *N*, as if the read behaved according to our assumptions.

When there is evidence that the target is unique, the mapping quality is commensurate with the belief that *N* = 0. In this case, it would be more appropriate to describe the estimation of *N* as measuring the probability that a paralog was missed during the search. This is the logic that explains the high mapping quality of super reads: when there is a perfect match, the screening process described in Figure 4 implicitly rules out many scenarios where the target has a paralog in the genome (Figure 5), so that only rare events remain possible. The calculations are completely independent of `sesame` and even independent of the seeding process, they are thus very general and can be used in any mapper based on the FM-index. In this sense, the introduction of super reads constitutes one of the most important contributions of the present work.

Figure 6 suggests that the mapping quality of super reads is properly calibrated but the representation is somewhat self serving because in reality the true error rate cannot be computed in this case. A mapping quality of 120 means one error every 2000 sequencing runs (assuming that all 500 million reads of each run are super reads), meaning that in practice one will never observe a mapping error for reads in this category. So the exact MAPQ score above 120 does not matter, as long as all the reads are mapped correctly. For super reads of size 50, the mapping quality is around 75 and it seems to be correctly estimated based on the simulated data from the human genome (sixth panel from the left in Figure 6), there is thus no reason to doubt the estimates at higher quality.

Mapping quality increases with the read length (Figure 6), but longer reads are less likely to be super. The reason is simply that they are more likely to contain a sequencing error. This appears on the rightmost panel of Figure 6, where in the pine genome, there are more reads of size 50 that map without mistakes than reads of size 100. This suggests that the strategy of MMP could be improved by running the uniqueness test of Figure 4 only on the parts of the reads that have a perfect match in the genome. This would allow more reads to be labelled as super, though with a lower mapping quality. But overall, future improvements are more likely to come from better indexing, as we explain below.

### 4.2 Performance

Figure 8 shows that MMP is 2–4 times faster than BWA-MEM and Bowtie2, while using 2–3 times more memory (trading speed for memory in MMP was a design decision based on the standards of present-day hardware). It is probable that BWA-MEM and Bowtie2 would also run 2–3 times faster with an equally large index size, but the strategy of not investing time in reads that cannot be mapped accurately seems valid. It is also important to bear in mind that our benchmark is not entirely fair because BWA-MEM and Bowtie2 are general mappers: they have to meet the demand on many different tasks so they cannot use some of the shortcuts implemeted in MMP.

Overall the mappers spend time on very different tasks: BWA-MEM and Bowtie2 use sophisticated methods to refine the candidate set after seeding, for instance by extending the set through re-seeding. In comparison MMP either tests all the candidates (when *N* ≤ 20) or tests only one (when *N >* 20), but it never extends the candidate set. The extra work that has to be done by MMP is to estimate *N* (Figure 3), test uniqueness (Figure 4) and compute mapping qualities with `sesame`. The running time of `sesame` is negligible, but the other two tasks represent a significant part of the running time (around 10–20% with large variations on different data sets). The point of MMP is to demonstrate that these steps can be carried out fast enough for a mapper to remain competitive.

However, it is wasteful to test uniqueness for every read. Unique sequences can be flagged at indexing time so that no additional computations are required at mapping time. For instance, if it is known with absolute certainty that a locus has no paralog, then mapping quality is extremely high throughout the locus (remember that spurious random hits are easy to detect as soon as the reads are longer than approximately 30 nucleotides). With this kind of information, super reads would be unnecessary and even higher mapping quality could be achieved on large parts of the genome. Likewise, the number of paralogs *N* of each locus could be stored to save computation time and to obtain more accurate estimates of the mapping quality throughout the genome, somewhat analogous to the concept of mappability [18]. It is presently unclear how to compute *N* for every locus and how to annotate a genome accordingly, but when practical solutions exist, faithful mapping will gain speed and robustness.

The case of accuracy is interesting (Figure 7). For the genome of *Drosophila*, which is relatively simple, the benefits of MMP are evident, while they are mitigated for the more complex human and pine genomes. MMP tends to “give up” easily to save time on difficult reads, so it may underperform when these cases are too frequent. One straightfoward way to improve the accuracy would be to map the reads with a more sensitive strategy when their mapping quality is too low. For instance, using skip seeds like Bowtie2 may redeem such reads, if one is willing to spend the time to give them a second chance. This would not be a problem because the seeding probabilities of skip seeds are readily available in `sesame`. The only downside would be a loss of speed, but the mapper would still be faithful.

### 4.3 Benefits of faithul mapping

The idea of developing a faithful heuristic for better calibration was originally proposed in BLAST [23]. The speed gain over its ancestor FASTP was very substantial, whereas this is not the case for MMP. Modern short-read mappers are highly optimized so it is unlikely that a 10-fold speedup could be gained by just calibrating the heuristics they rely on. That said, MMP demonstrates that there is room for improvement by exploring new ways to efficiently estimate faithfulness.

The present work focuses on general mapping tasks in eukaryotic genomes, but faithful read mappers may find much more interesting applications, such as when the reference contains several variants of the same sequences. This is the case for the genome of hybrid species or of heterozygote individuals where each sequence has *N* ≥ 1 paralogs in the reference. In this case, the challenge is to estimate the confidence that the read maps to one genome or the other. MMP could be used for this kind of problem, but a more specialized method that capitalizes on the key information *N* ≥ 1 would be more appropriate. For instance, the time spent looking for super reads is wasted in this context because they cannot occur (not a single sequence is unique). A more specialized method dedicated to calibrating low mapping quality is expected to give better results.

Another case of interest is when the DNA sample is contaminated and needs to be mapped to several species. This occurs when working with primates because experimenters can contaminate the biological material with their own DNA, or simply when the source of the DNA is unknown. In such cases, only few sequences are expected to have no paralog in the reference. In particular, if the genomes of interest are relatively close, the copy number will usually be equal to the number of genomes. In such cases, a general faithful mapper such as MMP is expected to give decent results, even though it was not tested in this context. More generally, we expect faithful mapping algorithms to find other applications in many areas of biology.

## 5 CONCLUSION

Faithfulness is an important feature of the mapping process. We have demonstrated that it is possible to achieve faithful mapping while remaining competitive in terms of speed, accuracy and memory usage. Exploring new algorithms for faithful mapping is a promising avenue of research that can also bring benefits in speed and mapping quality.

## 6 ACKNOWLEDGEMENTS

We would like to thank Santiago Marco-Sola, Catalina Romero and Nicholas Stroustrup for their critical comments on the manuscript. This work was funded by the Spanish Ministry of Economy, Industry and Competitiveness (MEIC, Plan Estatal PGC2018-099807-B-I00) and by the European Research Council (Synergy Grant 609989). R. C. was supported by the People Programme (Marie Curie Actions) of the European Union’s Seventh Framework Programme (FP7/2007-2013) under REA grant agreement 608959. We acknowledge the financial support of the MEIC to the EMBL partnership, the Centro de Excelencia Severo Ochoa and the CERCA Programme / Generalitat de Catalunya.

