## Supplementary Figures for "Mapping short reads, faithfully"

### Mapping short reads, faithfully: supplementary figures

February 10, 2020

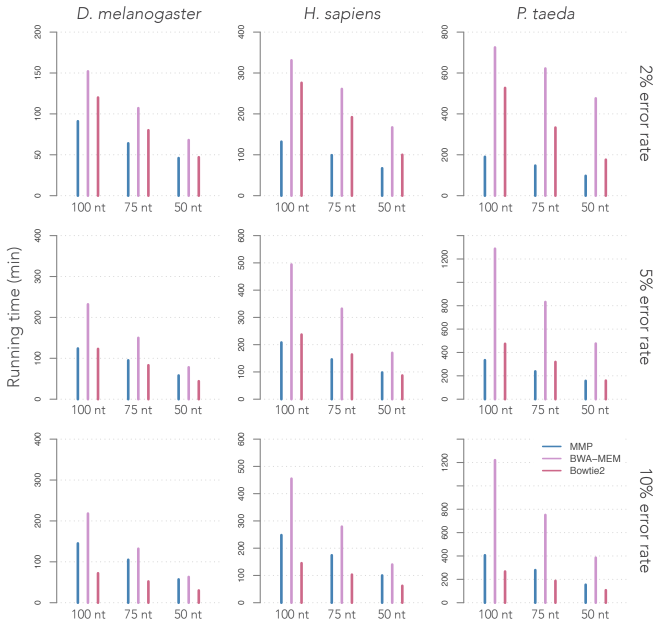


Supplementary Figure 1: Resource usage at 2, 5 and 10% error rate. The representations are as in Figure 8 (the memory footprint does not depend on the error rate of the sequencing process).

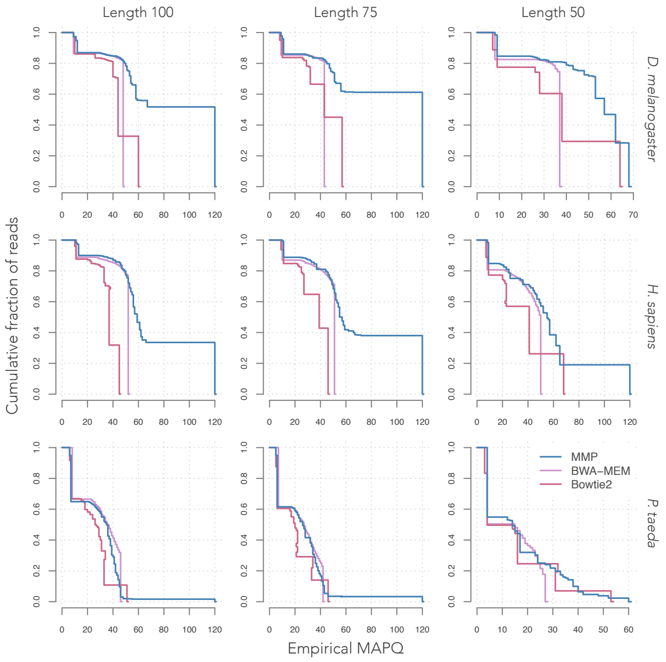


Supplementary Figure 2: Mapping accuracy with 2% error rate. The representations are as in Figure 7.

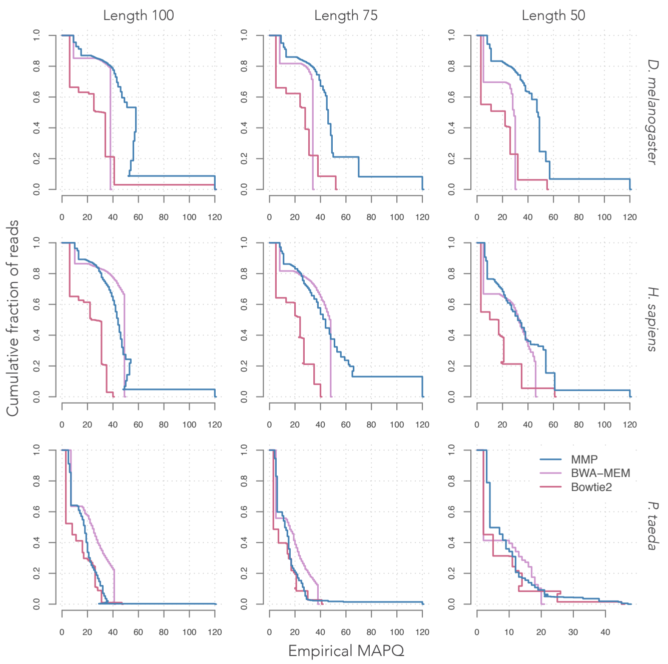


Supplementary Figure 3: Mapping accuracy with 5% error rate. The representations are as in Figure 7.

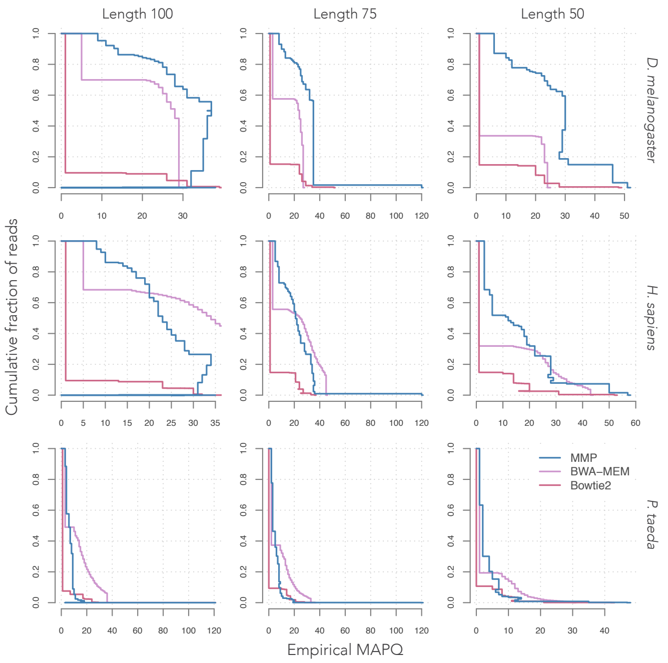


Supplementary Figure 4: Mapping accuracy with 10% error rate. The representations are as in Figure 7.

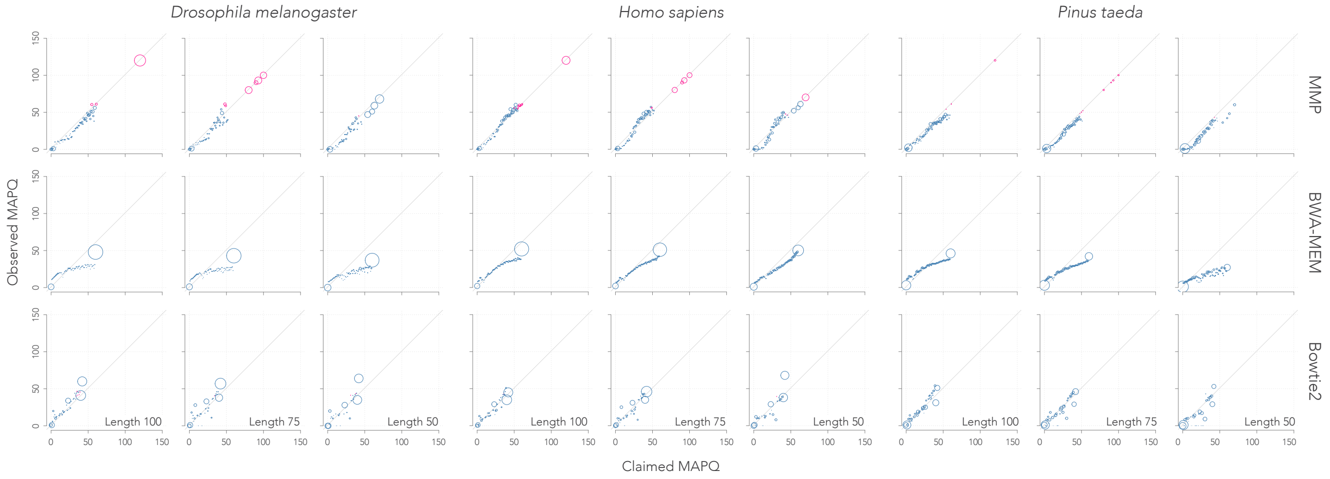


Supplementary Figure 5: Faithfulness with 2% error rate. The representations are the same as in Figure 6.

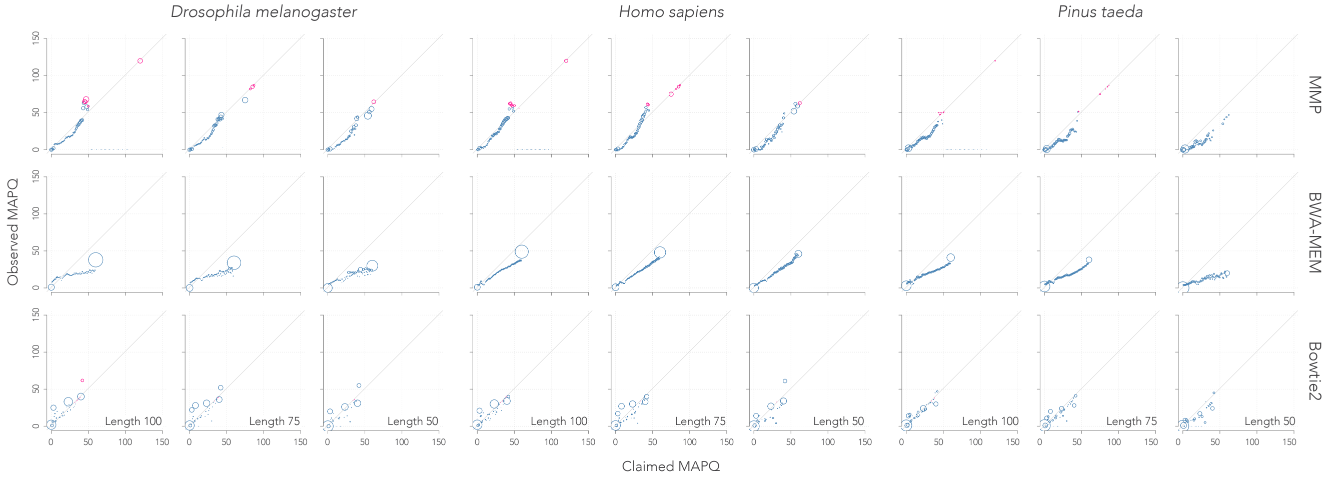


Supplementary Figure 6: Faithfulness with 5% error rate. The representations are the same as in Figure 6.

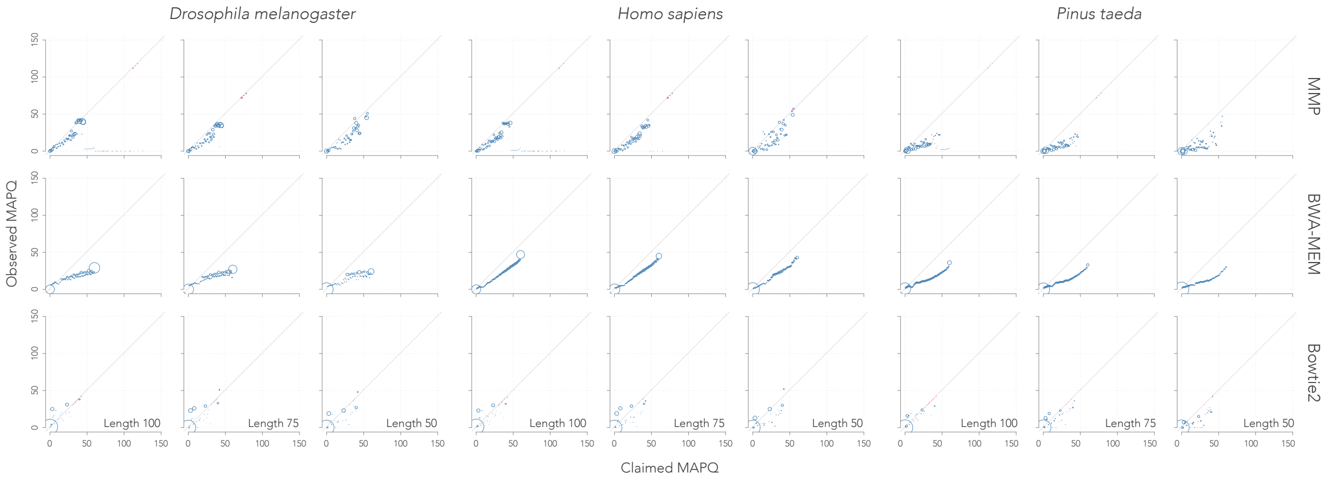


Supplementary Figure 7: Faithfulness with 10% error rate. The representations are the same as in Figure 6.
